# Towards a further understanding of measles vaccine hesitancy in Khartoum state, Sudan: A qualitative study

**DOI:** 10.1101/568345

**Authors:** Majdi M. Sabahelzain, Mohamed Moukhyer, Eve Dubé, Ahmed Hardan, Hans Bosma, Bart van den Borne

## Abstract

**Background:** Vaccine hesitancy is one of the contributors to low vaccination coverage in both developed and developing countries. Sudan is one of the countries that suffers from low measles vaccine coverage and from measles outbreaks. For a further understanding of measles vaccine hesitancy in Sudan, this study aimed at exploring the opinions of Expanded Program on Immunization officers at ministries of health, WHO, UNICEF and vaccine care providers at Khartoum-based primary healthcare centers.

**Methods:** Qualitative data were collected using semi-structured interviews during the period January-March 2018. The topic list for the interviews was developed and analyzed using the framework “Determinants of Vaccine Hesitancy Matrix” that developed by the WHO-SAGE Working Group.

**Findings:** The interviews were conducted with 14 participants. The majority of participants confirmed the existence of measles vaccine hesitancy in Khartoum state. They further identified various determinants that grouped into three domains including contextual, groups and vaccination influences. The main contextual determinant as reported is the presence of “anti-vaccination”; who mostly belong to some religious and ethnic groups. Parents’ beliefs about prevention and treatment from measles are the main determinants of the group influences. Attitude of the vaccine providers, measles vaccine schedule and its mode of delivery were the main vaccine related determinants.

**Conclusion:** Measles vaccine hesitancy in Sudan appears complex and highly specific to local circumstances. To better understand the magnitude and the context-specific causes of measles vaccine hesitancy and to develop adapted strategies to address them, there is a need to investigate measles vaccine hesitancy among parents.

## Introduction

Globally, vaccination is recognized as one of the most cost-effective public health measures [1]. It is estimated that a 60% reduction in measles mortality occurred worldwide since 1999, however, measles still remains a leading cause of vaccine-preventable death in children aged less than 5 years in sub-Saharan Africa. [2]

Over the past twenty years, concerns have been raised regarding the spurious link between the measles, mumps, rubella (MMR) vaccines and development of autism and autism spectrum disorders (ASD) [3]. Such concerns contributed to a decrease in vaccine uptake and an increase in the number of cases of many vaccine preventable diseases over the past several years in Europe and the USA [4–6].

Vaccine hesitancy is one of the contributors to a low vaccination coverage in both developed and developing countries [7]. As defined by the WHO Strategic Advisory Group on Experts (SAGE) on Immunization: “Vaccine hesitancy refers to a delay in acceptance or refusal of vaccination despite availability of vaccination services. Vaccine hesitancy is complex and context-specific, varying across time and place. It is influenced by factors, such as complacency, convenience, and confidence” [8]. Complacency exists where perceived risks of vaccine-preventable diseases are low; convenience relates to access issues such as the physical availability, ability to understand (language and health literacy), and vaccine confidence is defined as the level of trust in a vaccine or provider.[6]

In Sudan, the Expanded Program on Immunization (EPI) was launched in 1976. The EPI services are provided free of charge through the Primary Health Care centers. Communication strategies that address vaccine hesitancy were adopted as a policy by the federal ministry of health in order to enhance community demand for vaccines [9].

Sudan is still far from eliminating measles, as the measles vaccine coverage should be and sustained at 95% for the first (MCV1) and second (MCV2) doses of the vaccine, however, in 2017 the national coverage was 90% and 72% for MCV1 and MCV2 respectively [10]. Another indicator is to reduce the endemic incidence of measles to zero cases per 1 million of population. However, in 2017, an unpublished report from WHO indicated that the incidence of measles was 14.2 per 1 million.

Little is known about the determinants of the low measles vaccine coverage in Sudan. For a further understanding of measles vaccine hesitancy in Sudan, this study aims to explore the opinions of Expanded Program on Immunization (EPI) officers/experts and front-line vaccine providers who are based in Khartoum state.

## Methods

### Study design

This study had a cross-sectional design and data were collected using semi-structured interviews during the period January - March 2018. The topic list for the interviews was developed and structured using the model framework “Determinants of Vaccine Hesitancy Matrix” that was developed by the WHO-SAGE Working Group. It distinguishes between three groups of determinants: contextual influences, such as religion and geographic barriers, individual and group influences, such as beliefs and attitudes about health and prevention, and vaccine or vaccination-specific influences, such as the mode of administration and the vaccination schedule.

### Groups and participants selection

Two groups were included for the individual interviews. Firstly, the Expanded Program on Immunization (EPI) officers/ experts at the federal ministry of health, Khartoum state ministry of health, WHO and UNICEF. The EPI officers/experts were selected for having at least ten years of experience in the field of immunization and having a leading role inside and/or outside Khartoum state regarding the immunization program. Secondly, front-line vaccine providers at primary healthcare centers. The vaccine providers were selected purposively from areas with low measles vaccine coverage or areas that had previous measles outbreaks in rural and urban areas in Khartoum state. Taking into consideration the administrative subdivision of Khartoum state into seven localities, the vaccine providers were selected from all localities, which have urban and rural areas, except for one locality with only an urban area. This diversity in the two groups enabled us to understand in-depth insight into the salience of measles vaccine hesitancy among Sudanese parents from the perspective of experts and providers.

### Interview questions

Questions about the causes and determinants of vaccine hesitancy-measles vaccine hesitancy were explained to the participants. Both groups were asked questions about the existence and determinants of vaccine hesitancy in Khartoum state. Some questions were asked only to the group of EPI officers/experts, as these questions required long experience and expertise at the national and Khartoum state levels, such as their opinions about the definition of vaccine hesitancy and its impact on the overall measles vaccine coverage.

### Study area

This study was conducted in Khartoum state, where vaccination services are comparatively easily available. This allows the study of hesitancy. Khartoum state also inhabits a diversity of groups of people in terms of socio-cultural and socio-economic backgrounds and exposure to vaccination campaigns and materials. Khartoum state has an area of 22,122 km^2^ and an estimated population of approximately 7,152,102 (2008). Administratively, Khartoum State is divided into seven localities, namely, Khartoum, Jabel Awleya, Khartoum North and East Nile, Omdurman, Umbada and Karari.

### Data collection and analysis

The individual interviews with the participants from the two groups were conducted in the Arabic language. The data were recorded, transcribed, coded manually, and then analyzed according to the model framework of the WHO-SAGE of determinants of vaccine hesitancy by the principle investigator (MS); then another researcher (ED) reviewed the work. Hence, the themes and sub-themes were also clustered according to the model framework. Afterwards, expressive quotes were selected from the transcription, translated into English, and presented in the text. The translation was checked by a senior researcher.

### Ethical Consideration

The study was approved by the Ahfad University for women’s Review Board (IRB) and the National Health Research Ethics Committee at the Federal Ministry of Health in Sudan (Proposal No.1-1-2018).

## Results

The interviews were completed with 14 participants, as saturation was reached after 13 interviews. Five were EPI officers/experts and nine were front-line vaccine providers. The demographic and professional characteristics of the participants are summarized in Table 1. Three themes emerged from the interviews: definition of vaccine hesitancy, determinants of vaccine hesitancy (contextual, groups and vaccine related determinants) and the effect of vaccine hesitancy on measles vaccine coverage.

**Table 1.** The Demographic and professional characteristics of the participants.

| No. | Participants | Sex | Age | Experience<br>(Years) | Qualification | Profession and Place of<br>Work |
| --- | --- | --- | --- | --- | --- | --- |
| 1 | A | M | 47 | 16 | - | EPI officers/Expert |
| 2 | B | M | - | - | - | EPI officers/Expert |
| 3 | C | M | 50 | 10 | - | EPI officers/Expert |
| 4 | D | F | 42 | 15 | - | EPI officers/Expert |
| 5 | E | M | - | 15 | - | EPI officers/Expert |
| 6 | Fp | F | 36 | 10 | - | VP, Urban area |
| 7 | Gp | M | 58 | 21 | Medium School | VP, Rural area |
| 8 | Hp | F | 45 | 20 | Secondary School | VP, Urban area |
| 9 | Ip | F | 40 | 15 | Secondary School | VP, Urban area |
| 10 | Jp | F | 49 | 21 | Secondary School | VP, Rural area |
| 11 | Kp | F | 41 | 11 | Secondary School | VP, Rural area |
| 12 | Lp | F | - | - | - | VP, Urban area |
| 13 | Mp | F | - | - | - | VP, Urban area |
| 14 | Np | F | 38 | 17 | Secondary School | VP, Rural area |

### Definition of vaccine hesitancy from the perspectives of the EPI officer/Experts

The five EPI officers/experts were asked how they would define vaccine hesitancy. Several answers were reported; one participant defined it as inability of a person to access and utilize the vaccines services (participant C). Two participants (Participants A and D) indicated that vaccine hesitancy is related to those who do not vaccinate at all “The hesitancy is present among people with a zero dose”. However, participant D added delaying of vaccination as a result of different concerns. Two participants associated vaccine hesitancy with “rumors” (Participant B and E)

> “The issue is that some people do not come to the vaccination in general, or delay vaccination for certain concerns; either religious, vaccine safety, or related to the service provided”

### Determinants of vaccine hesitancy in Khartoum state

Both groups of participants were asked about the main causes of vaccine hesitancy in Khartoum state. Fitting in the WHO Model of determinants of vaccine hesitancy, the findings are summarized in (Fig 1).

**Fig 1.**
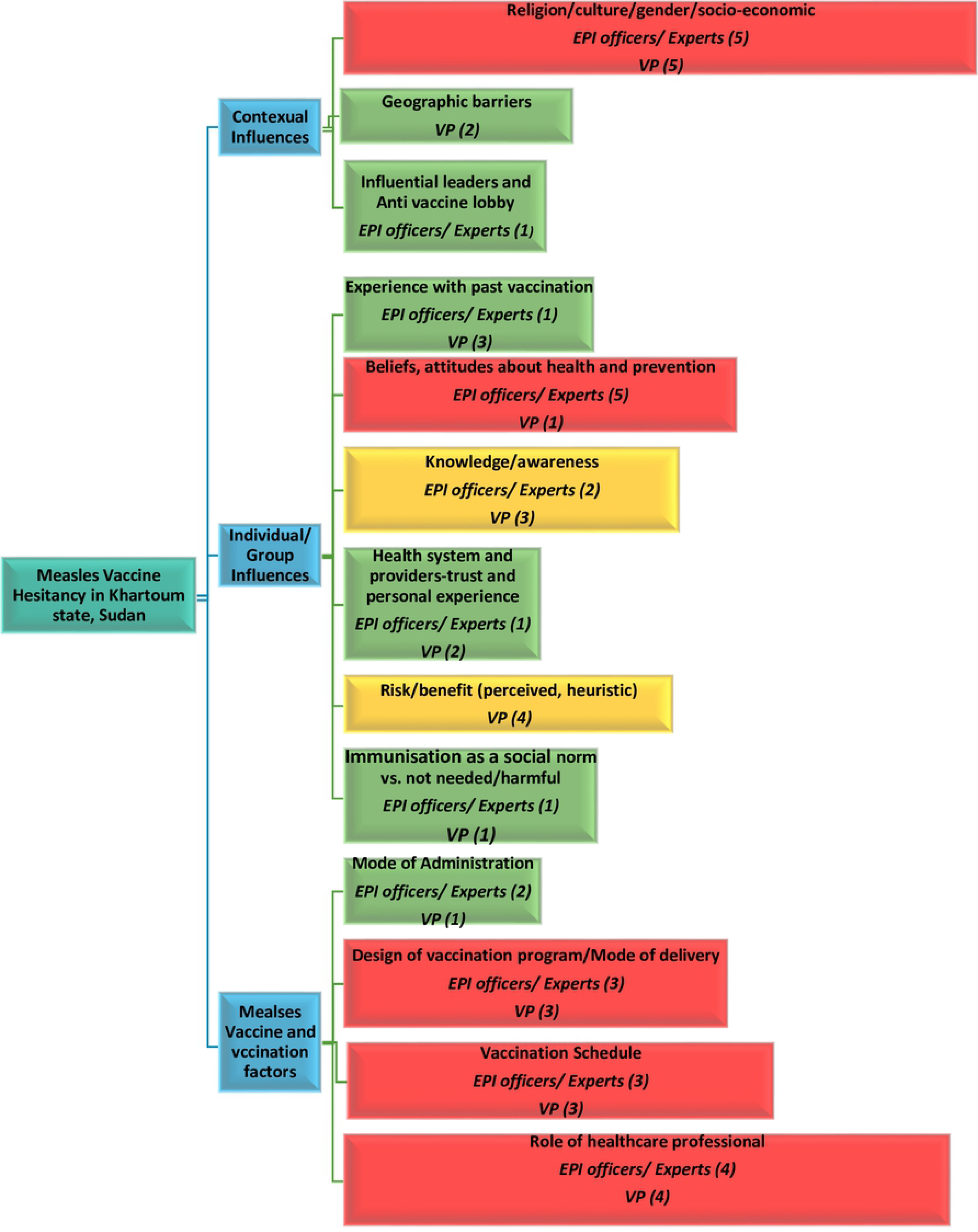
EPI officers/experts’ and Vaccines Providers’ (VP) opinions about determinants of measles vaccine hesitancy in Khartoum state, Sudan.

#### 1. Contextual Influences

Religious belief from certain religious groups were reported by the majority of the participants in both groups. However, some EPI officers/experts elaborated more on these beliefs in religious groups.

> “There is an escalation of religious refusal, for instance; groups such as Ansar Al-Sunna and Wahabia (Religious groups) think that vaccines are brought by Jews, for which reason, they refuse all vaccines”
>
> — (Participant C).

> “ Not only measles, but also all vaccines, sometimes they talk about Freemasons and infidel states … etc. which bring vaccines”
>
> — (Participant B)

> “A few years ago, some public figures adopted vaccine refusal. If you heard about Shaikh Sadig Abdallah Abdelmagid, the leader of the Muslim Brotherhood (religious and political group), who advocated the prohibition of child vaccination”
>
> — (Participant A).

Other groups in which vaccine hesitancy was identified by several immunization officers/experts included ethnic and tribal groups, people of higher socioeconomic status and well-educated people who live in first class areas in Khartoum state.

> “Some tribes such as Falatta (a tribe of a Nigerian origin) do not vaccinate all their children in their households; they are afraid of the evil eye (i.e. they think that people will notice that they have many children in their house, so they think some of them will die)”
>
> — (Participant C)

Few of the interviewed front-line providers noted geographic barrier as a cause of vaccine hesitancy, however, it was considered a minor issue by some immunization officers/experts.

> “The higher population density is to the west of the asphalt street, but the Health Center is to the east of the asphalt street, so mothers suffer when they bring children, walking all this distance”
>
> — (Participant N*p*).

> “ Reaching the health centers is not an issue, we have village clinics in all of Khartoum State, and we have mobile teams in some localities such as Umbada and Jebel Awlya, even in the Internal Displaced People camps”
>
> — (Participant A).

#### 2. Individual and group influences

With consensus from all (five) immunization officers/experts, beliefs and attitudes of parents/guardians about health and prevention from measles was identified as a causal factor in measles vaccine hesitancy. It is interesting to note that only one participant from front-line providers reported beliefs and attitude of parents/guardians as a contributing factor.

> “Measles is believed to be a common household disease that every child contracts, with mild fever and mild rash, then topical medication or anything else is used, and then the disease disappears”
>
> — (Participant D)

> “Ansar Al-Sunna (i.e people who belong to this group) say it is god who protects us until we grew up, vaccination didn’t exist in the past”
>
> — (Participant F*p*).

Experience with past vaccination was identified by some of the front-line providers and mentioned by only one immunization officer/expert as a possible cause.

> “ When the child cries at night due to fever after vaccination, the father tells his wife that she took a healthy child, who is now ill, telling the mother not to take the child again for vaccination”
>
> — (Participant K*p*)

Some participants from both groups reported lack of knowledge and awareness as an important determinant in measles vaccine hesitancy.

> “ There is a lack of knowledgeable among mothers, I mean mothers’ awareness about coming at the exact time of the measles vaccination, or the importance of vaccinating with measles vaccine, similarly when the time comes for the second dose at age 18 months”
>
> — (Participant D)

Many front-line providers noted that perception of a lack of risk and low benefit of vaccination among people (parents/guardians) are important determinants of measles vaccine hesitancy.

> “Measles is rare nowadays, not as used to be in the past, when measles caused death. Now some mothers believe that there is no need for this vaccine”
>
> — (Participant F*p*).

Immunization is not considered as a social norm by some people in Sudan; this was reported by one of the front-line provider

> “A woman came to me and said that she is a nomad from Western Sudan, she said that for us cattle are the most important (i.e. it is not our priority)”
>
> — (Participant I*p*).

#### 3. Vaccine- and vaccination-specific factors

Mode of administration was identified as a contributing factor in measles vaccine hesitancy by two immunization officers/experts and one front-line providers.

> “when a mother feels that her child is going strong at the age of 9 months, they think that this child is well and that there is no need to vaccinate him or to expose him to an injection (i.e. measles vaccine)”
>
> — (Participant C).

Some of the front line providers noted that the design of the vaccination program or the mode of delivery as an important factors in measles vaccine hesitancy, specifically the number of measles vaccination’s sessions per week (As most of the vaccine providers indicated that they have one session per week for measles vaccination in order to comply with the recommendation of discarding the ten-dose vial after six hours from opening the vial as well as reducing the loss of unused doses). However, some of the immunization officers/ experts thought this to be of less influence.

> “ One day, three women came to me and went back, I told them that the measles vaccine was not that day, measles vaccination is on Wednesdays. They got upset, because it was hard to come again”
>
> — (Participant J*p*).

The majority of participants in both groups identified the measles vaccine schedule (Measles vaccine is given at ninth and eighteenth months of child age. Vaccine providers use different ways for reminding parents about the time of the next dose including telling verbally, write on the card, and/ or even using mobile phone call.) as a causal factor in measles vaccine hesitancy.

> “.. the long gap between scheduled vaccinations means that the mothers might forget, though we remind them by writing in their cards the exact time of the following dose”
>
> — (Participant L*p*)

The attitude and the personal characteristic of vaccine providers was reported by most of the participants in the both groups as a contributing factor in vaccine hesitancy (e.g. some parents especially in rural areas do not trust foreigners’ vaccine providers; prefer female providers..etc)

> “ Sometimes the cadre may be rough or is not from the same rural area. If such a vaccine provider and I are working together in the office, parents will ask me personally to vaccinate their children, and when I reply that he and I are the same, they say that they have no trust in someone from outside their village”
>
> — (Participant N*p*)

### The effect of Vaccine Hesitancy on measles vaccine coverage

The immunization officers/ experts were asked a question about the effect of vaccine hesitancy on measles vaccine coverage in Khartoum state. Some of them reported it as a causal factor in reducing measles vaccine coverage, however, fewer considered it as a contributing factor with manty other challenges including community awareness about the importance of vaccination.

> “ There are several reasons for reduced coverage, but I still say that it is of great importance to stress the necessity for both the pentavalent vaccine and the measles vaccine, according to the recommended schedules”
>
> — (Participant B).

## Discussion

For a further understanding of measles vaccine hesitancy in Khartoum state, Sudan, this study explored the views of EPI officers/experts and front-line vaccine providers. The findings show that there was no general consensus on the definition of vaccine hesitancy among the EPI officers/experts, nor was there a strong consistency with the definition of the WHO-SAGE group. Some limited their definition to the refusal of the vaccine. Findings in general confirm what other similar studies reported. [11, 12]

Despite this lack of consensus on the definition per se, our study’s findings indicate that the majority of participants, both EPI officers/experts and front-line vaccine providers, confirmed the existence of some form of ‘measles vaccine hesitancy’ in Khartoum state. Further, they agreed on it as a contributing factor for the decreasing measles vaccine coverage in Khartoum and likely in all of Sudan.

Most of the participants in both groups reported many contextual, individual/group and vaccination/vaccine issues as determinants of measles vaccine hesitancy. The majority of the participants agreed that the main contextual determinant is the presence of people (parents/guardians) who can be qualified as “anti-vaccination”; they mostly belong to religious groups, especially the Ansar Al-Sunna group, and they often refuse all vaccines. Some of the EPI officers/experts attributed this refusal to blaming the manufacturing countries of a conspiracy.

Other vaccine providers explained the refusal by pointing to particular beliefs, such as the belief that the vaccines are a cause of the disease and the belief that only God can protect their children from disease. These findings are congruent with many studies that linked vaccine hesitancy with certain religious, ethnic and cultural groups in different countries [12-15]. Our study’s findings reflect how these different beliefs concerning vaccination are rooted in particular sub-groups.

Appropriate interventions are needed to address these different beliefs in order to increase the measles vaccine coverage.

Moreover, most EPI officer/experts agreed that individual beliefs and attitudes about health and prevention from measles are the main determinants (causal factors) of measles vaccine hesitancy, and they could identify some of these beliefs. This is consistent with findings from France, where one of the major reasons for non-vaccination is the perception that the risk of acquiring measles is low [16]. None of the participants in our study reported concerns about a possible link between measles vaccination and autism.

Most participants agreed about three factors that relate to vaccination issues and vaccine hesitancy in Sudan. These factors include personal characteristics and the attitudes of the vaccine providers, the measles vaccination schedule (i.e. the Ninth month’ and the Eighteen month’s doses), and the mode of measles vaccine delivery (e.g. limiting the number of opening sessions for measles vaccination per week). As most of the vaccine providers indicated that, they have one session per week for measles vaccination in order to comply with the recommendation of discarding the ten-dose vial after six hours from opening the vial as well as reducing the loss of unused doses. These measles vaccine and vaccination-related issues lead to missing the opportunity for vaccination; parents come to vaccinate their children and find the measles vaccination’s session at another day, for which they might not return.

### Limitations of the study

Firstly, this study was qualitative and Khartoum-based. Furthermore, the selection of vaccine providers was also intentionally biased as it was done purposively in geographical areas where measles vaccination uptake was low or where measles outbreaks were reported to be more frequent and prevalent. Hence, findings cannot be generalized to the whole population of vaccine providers and EPI officers/experts in Sudan. Secondly, most participants mentioned names of some ethnic and religious groups without a clear definition of who exactly these groups are (e.g. “*Falatta* “ and “Ansar Al-Sunna”). This might complicate the search for the right target group for interventions. We did this study into vaccine hesitancy in circumstances where people have a relatively easy access to vaccination services. We have to acknowledge that there are other areas in Sudan, where there is the perhaps even more fundamental difficulty of people not having access to vaccination services in the first place.

## Conclusion

To conclude, the findings of this study show how complex measles hesitancy is in Sudan and how much variation there is regarding the definition and the perceived causes and consequences. It also indicates that this risky phenomenon can only be explained (and thus possibly be intervened upon) in its specific socioeconomic and –cultural context. Negative beliefs and attitudes of people (parents/guardians) are important, but vaccination program aspects should not be neglected. For an even further understanding of the magnitude of the problem and the context-specific causes of measles vaccine hesitancy and for developing better intervention strategies to address hesitancy, there is a clear need to study the issue in even more detail, including among the parents/guardians themselves.

## Acknowledgements

We would like to thank the participants from the two groups. We appreciate many government officials at federal and state levels for their cooperation and providing necessary information and permissions to conduct the study. Special words and thanks to Dr. Awad Hassan from Ahfad University for reviewing English translation of the quotes, and to Dr. Howida Hassan, Sara Said, Fatin Hashim, Fatima Suliman and Mozdalifa Mohamed for their help in transcribing the data.

## Supporting information

S1 File. Questions guide

